# Pharmacological Studies on the Efficacy of Polyherbal Formulation Dawa ul kurkum in DEN-Induced 2-AAF-Promoted Hepatocellular Carcinoma in Male Wistar Rats

**DOI:** 10.1101/2023.05.05.539655

**Authors:** Meenakshi Gupta, Hemlata Nimesh, Maryam Sarwat

## Abstract

The purpose of this study was to explore the anti-cancer activity and the possible mechanism of Dawa-ul kurkum (Duk) against diethylnitrosamine (DEN)-initiated hepatocarcinogenesis in male Wistar rats. We administered Duk at 3 doses, viz., 75, 150, and 300 mg/kg/day, 2 weeks before the DEN and continued it for 16 weeks. We found that Duk significantly reduced the DEN and 2-AAF induced phenotypical changes in rats and restored the activities of serum markers. Furthermore, Duk counteracted the oxidative stress induced by carcinogens as observed by restoration in the levels of superoxide dismutase (SOD) and catalase (CAT). Duk significantly diminished the levels of malondialdehyde (MDA) in a dose dependent manner and restored the liver microarchitecture as assessed by histopathological studies. The results of immunohistochemical staining showed that Duk inhibited the DEN-induced decrease in the number of cells positive for Bid and Caspase-9. It also reduces the number of cells positive for Cyclin D. This study shows that Duk significantly protects rat liver from hepatocarcinogenesis by regulating oxidative damage and restoring serum markers. The chemopreventive effect of Duk might be through induction of apoptosis.

## 1. Introduction

Hepatocellular carcinoma (HCC), the sixth most common progressively increasing solid tumor, is also the second major cause of cancer mortality worldwide. It is a chemo-resistant tumor, causing the death of about 80% of patients within a year of diagnosis [1]. Although various treatment strategies such as radiotherapy, surgical resection, embolization, ablation, and chemotherapy are available, they all have limitations. They are known to cause broad-spectrum adverse effects and toxicities leading to poor selectivity toward tumor tissues and multi-drug resistance against chemotherapeutic agents. Management with sorafenib shows survival benefits for 9-11 months, still leaving the relevant population without effective therapy [2-4]. Therefore, an increasing number of patients and even oncologists are looking for alternative approaches to manage HCC.

Complementary and alternative medicine (CAM) approaches can be used to improve the quality of life and may prolong the survival of patients [5]. World Health Organization has recognized Unani medical system as efficient alternative medicine systems [6]. It is very well practiced in China, Egypt, India, Iraq, Persia, and Syria. Thus this approach has become very credible. Traditional medicine practitioners like Ayurveda, Amchi and Unani believe that the treatment of complex diseases like cancer can be addressed better with polyherbal mixtures than with a single medicinal plant [7]. This is because polyherbal preparations offer synergistic effect, better therapeutic outcome, and cause less toxicity [8]. One of the popular polyherbal Unani formulation is Dawa-ul-Kurkum (Duk). It is very popular among Unani practitioners for the treatment of liver dysfunction, anorexia, ascites, and abdominal pain [9,10]. However, its role for the treatment of HCC was not known. So, we designed this study to see the effect of Duk for the treatment of HCC in Wistar rats.

Duk consists of seven herbal components *Crocus sativus* L., *Nardostachys jatamansi* (D.Don) DC., *Cinnamomum cassia* (L.) J. Presl, *Cymbopogon jwarancusa* (Jones ex Roxb.) Schult., *Commiphora wightii* (Arn.) Bhandari, *Saussurea lappa* (Decne.) Sch.Bip. and *Cinnamomum zeylanicum* Blume (Figure S1). Literature shows anti-oxidant, an-ti-inflammatory, and anti-cancer properties of its constituent medicinal plants [11-17]. Various active components of its constituent medicinal plants have also reflected hepatoprotective activity [18-19].

Our previous results indicated the presence of various phytochemicals, viz., flavonoids, phenols, quinones, glycosides, etc. in Duk. It showed excellent antioxidant potential through DPPH and FRAP assay. We have also done the phytochemical characterization of Duk with the principal natural compounds of all the seven individual herbal components of Duk [10]. In the present study we have investigated the protective effect of Duk in diethylnitrosamine (DEN)-induced 2-aminoacetylfluorine (2-AAF)-promoted hepatocellular carcinoma in male Wistar rats. Using this well described model, we evaluated the effect of Duk on the serum markers, oxidative stress parameters and histopathological alterations in HCC-challenged rats.

## 2. Materials and Methods

### 2.1. Plant material, authentication and preparation of Duk

Duk was prepared from all of its seven herbal components by grounding them to powder and mixing them. The mixture was subjected to ethanol extraction (70% ethanol v/v) and filtration. The filtrate was evaporated, freeze dried, and stored in dark coloured bottles at 4 ºC [10].

### 2.2. Animals

A total of 48 Male Wistar Rats (100-125 g) procured from the animal house facility of Amity University, Noida, kept in clean cages (22 ± 2.4 ºC), fed with Harlan Tekla 8640 food (Madison, WI, USA) and filtered water ad libitum. During the experimentation, rats were kept under the normal standard light and dark cycle. All the experimental procedures of this study were duly ratified by the Institutional Animal Care and Ethical Committee of Amity University, Noida, Uttar Pradesh, India (CPCSEA/IAEC/AIP/2018/05/06). The recommendations cited by the Committee for Control and Supervision of Experiments and Animals (CPCSEA) were used as guidelines for maintaining the animals during this period. Rats were weighed on the first day of the experimentation and then every week.

### 2.3. Model and experimental design

The experimental HCC was initiated by DEN (Sigma Aldrich Co. LLC., St. Louis, MO, USA) and promoted by 2-AAF (Sigma Aldrich Co. LLC., St. Louis, MO, USA) [20]. Rats were kept on fasting for 4 days and were re-fed for 24 hours as mitotic proliferative stimulation. The experimental design and treatment protocol (Figure 1) were as follows:

**Figure 1.**
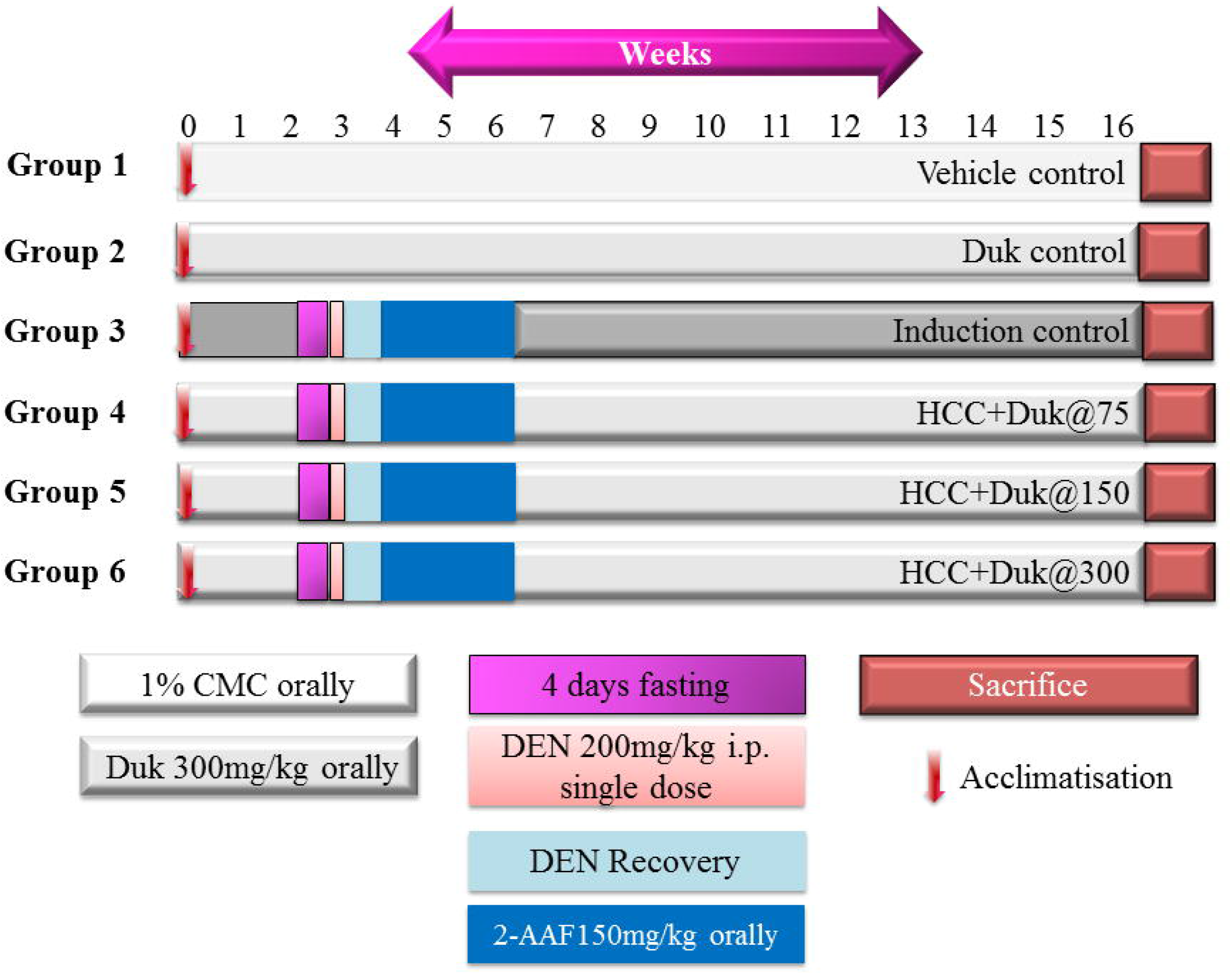
Details of experimental design. Male Wistar rats were divided into 6 groups (n=8). Group 1 (vehicle control; received 1% CMC, orally), Group 2 (Duk control; received Duk@300 mg/kg b.w., orally), Group 3 (Induction control, received DEN@200 mg/kg b.w., i.p. + 2-AAF@150 mg/kg b.w. orally), Group 4 (received carcinogens as group 3 + Duk@75 mg/kg b.w., orally), Group 5 (received carcinogens as group 3 + Duk@150 mg/kg b.w., orally), Group 6 (received carcinogens as group 3 + Duk@300 mg/kg b.w., orally). Duk was administered throughout the experiment and animals were sacrificed at the end of 16 weeks.

Group I: Normal control; animals were administered 1% carboxymethyl cellulose (CMC) orally for 120 days.

Group II: Duk Control; animals were orally administered Duk (300 mg/kg in 1% CMC) for 120 days

Group III: A single dose of DEN (200 mg/kg b.w.) was administered intraperitoneally. After 1 week of recovery, repeated doses of 2-AAF (150 mg/kg b.w.) were given orally for 12 days alternatively and animals were sacrificed at the end of 120 days.

Group IV: The same dose was administered as group III, along with 75 mg/kg Duk for 120 days.

Group V: The same dose was administered as group III, along with 150 mg/kg Duk for 120 days.

Group VI: The same dose was administered as group III, along with 300 mg/kg Duk for 120 days.

### 2.4. Tissue Preparation

At the end of the experiment period, all animals were anesthetized and the blood was collected. Serum was isolated and stored at −80 °C till further analyses. Animals were then sacrificed and livers were removed immediately and flooded with ice-cold saline. Homogenization was then carried out (at 4 °C) in 0.1 M phosphate buffer (pH 7.4) containing protease inhibitors (Genetix BioAsia Pvt. Ltd., Delhi, India). The homogenate was centrifuged (@800 g, 4 °C, 5 min) to separate nuclear debris and supernatant was used to estimate thiobarbituric reactive substances. For other biochemical assays, post-mitochondrial supernatant (PMS) was isolated by centrifuging the supernatant again (@10000 g, 4 °C, 20 min) [20].

### 2.5. Estimation of serum markers

The serum markers, aspartate aminotransferase (AST), alanine aminotransferase (ALT), and alkaline phosphatase (ALP), were measured spectrophotometrically. The assays were performed using commercially available standard enzymatic kits from Abcam Biotechnology Company (Cambridge, United Kingdom).

### 2.6. Estimation of oxidative stress markers

#### 2.6.1. Estimation of Catalase (CAT) levels

The CAT enzyme levels were determined by UV spectrophotometry. Briefly, 0.05 M phosphate buffer (pH 7), 0.019 M H2O2 and 50 μl PMS (in a total volume of 3 ml) were taken and the absorbance was recorded at 240 nm. CAT activity was expressed as na-nomoles H2O2 consumed per minute per milligram protein [20].

#### 2.6.2. Estimation of malondialdehyde (MDA) levels

100 μl PMS was mixed with 1.5 ml acetic acid (20%), 1.5 ml TBA (0.8%) and 200 μl sodium dodecyl sulfate (SDS; 8.1%) and the mixture was heated at 100 °C for 1 h. The mixture was then cooled and 5ml of n-butanol: pyridine (15:1%,v/v) and 1ml distilled water was added. The mixture was shaken vigorously and centrifuged (@4000 rpm, 10 min). The organic layer was separated and the MDA level was analyzed by measuring the absorbance at 532 nm using a spectrophotometer. The total amount of MDA formed in each sample was expressed as nanomoles MDA formed per milligram protein using a molar extinction coefficient of 1.56 × 10^5^ M^−1^ cm^−1^ [20].

#### 2.6.3. Estimation of Superoxide dismutase (SOD)

200 μl PMS was added to 800 μl of 50 mmol/l glycine buffer (pH 10.4) and 20 μl of a 20 mg/ml solution of (−) epinephrine. The absorbance was measured at 480 nm using a spectrophotometer. SOD activity was expressed as nanomoles of (−) epinephrine protected from oxidation by the sample using the molar extinction coefficient of 4.02×10^3^ M^−1^ cm^−1^ [20].

### 2.7. Histological examinations

For histological examinations, liver sections from different groups were stained with hematoxylin and eosin (H&E) [20]. Briefly, at the end of experiment, livers were removed and post-fixed in buffered formalin (10%) for 24 h. After fixation, slices (3–4 mm) of these tissues were dehydrated and embedded in paraffin. At least four cross-sections were taken from each tissue (5 μm thickness) and stained with H&E. Following two washings with xylene (2 min each), tissue sections were mounted with DPX mountant. The slides were observed for histopathological changes, and microphotographs were taken at ×20, ×40, and x60 magnifications.

### 2.8. Immunohistochemistry

For the immunohistochemistry experiments, blocks were de-paraffinized with three washes of xylene for 5 min each. The sections were then dehydrated with ascending alcohol concentrations and washed with distilled water (dH2O) two times for 5 min each. Sections were then immersed in pre-heated antigen retrieval solution, allowed to cool for 30 min, and then washed with dH_2_O three times for 5 min each. To reduce the nonspecific background staining, slides were incubated with 3% H_2_O_2_ for 10 min and washed with dH_2_O two times for 5 min each. Sections were washed in wash buffer (1X Tris Buffered Saline with Tween 20) for 5 min. After pre-treatment, the sections were incubated over-night at 4 °C with desired primary antibody (cat. no. #9662, #55506; Cell Signaling Tech-nologies, Inc. and PA5-29159; Thermo Fisher Scientific, Inc.) at 1:1000 dilutions. Following primary antibody treatment, sections were incubated (for 1 h) at room temperature with SignalStain® Boost Detection Reagent (HRP, cat. no. #8114; Cell Signaling Technologies, Inc.). Sections were washed three times with wash buffer and developed with freshly prepared and filtered 3,3’-diaminobenzidine (DAB) chromogen solution [21]. Sections were counterstained with hematoxylin, cleared, mounted, and visualized at a magnification of ×20, ×40, and x60.

### 2.9. Statistical analysis

Data were expressed as the mean ± SD. Data were analyzed using Graph Pad version 8.4.3 (San Diego, CA, USA) and compared using t-test and ANOVA. Differences between control and treatment groups were determined at a significance level, p>0.05.

## 3. Results

### 3.1. Duk restrains phenotypical changes in HCC rats

Treatment with DEN and 2-AAF is known to initiate cancer within 1 month. Varieties of modifications were observed; these included hair loss, reduction in the amount of food intake, and significant weight loss in the induction control animals as compared to control animals; all animals survived during the experimental period. These changes were restrained in animals treated with Duk in a dose dependent manner.

### 3.2. Duk restores serum markers and hepatic oxidative stress parameters

The serum activities of AST, ALP and ALT in induction control animals were considerably increased after DEN and 2-AAF administration and showed a clear correlation with hepatic dysfunction and HCC development. However, treatment with Duk showed a significant (p<0.05) and dose dependent fall in the levels of these serum markers (Figure 2).

**Figure 2.**
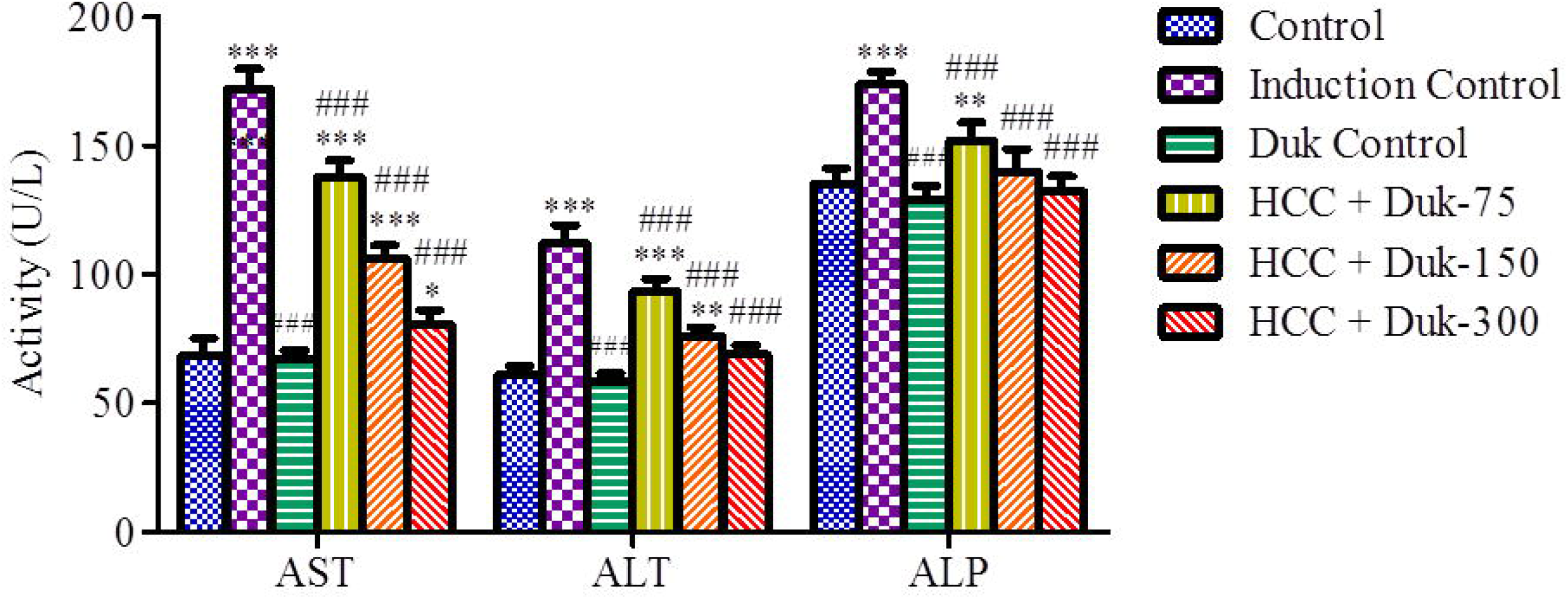
Level of AST, ALT and ALP in the serum of animals of all experimental groups. Values are expressed as Mean ± SD (P<0.05 vs. control group*; P<0.05 vs. induction control group#). Abbreviations: HCC: Hepatocellular Carcinoma; Duk: Dawa-ul-kurkum; HCC+Duk@75: Same dose of carcinogens as administered in induction control group + Duk at 75mg/kg dose; HCC+Duk@150: Same dose of carcinogens as administered in induction control group + Duk at 150mg/kg dose; HCC+Duk@300: Same dose of carcinogens as administered in induction control group + Duk at 300mg/kg dose; AST: Aspartate aminotransferase; ALT: Alanine transaminase; ALP: Alkaline phosphatase

Rats in the induction control group showed a significant increase in the levels of MDA and decrease in SOD and CAT activities after the administration of DEN+2-AAF, as compared to control group (Figure 3). However, treatment with Duk alleviated the levels the MDA in a dose dependent manner and brought significant increase in the levels of SOD and CAT. This indicated that Duk may attenuate the oxidative injury in HCC bearing rats.

**Figure 3.**
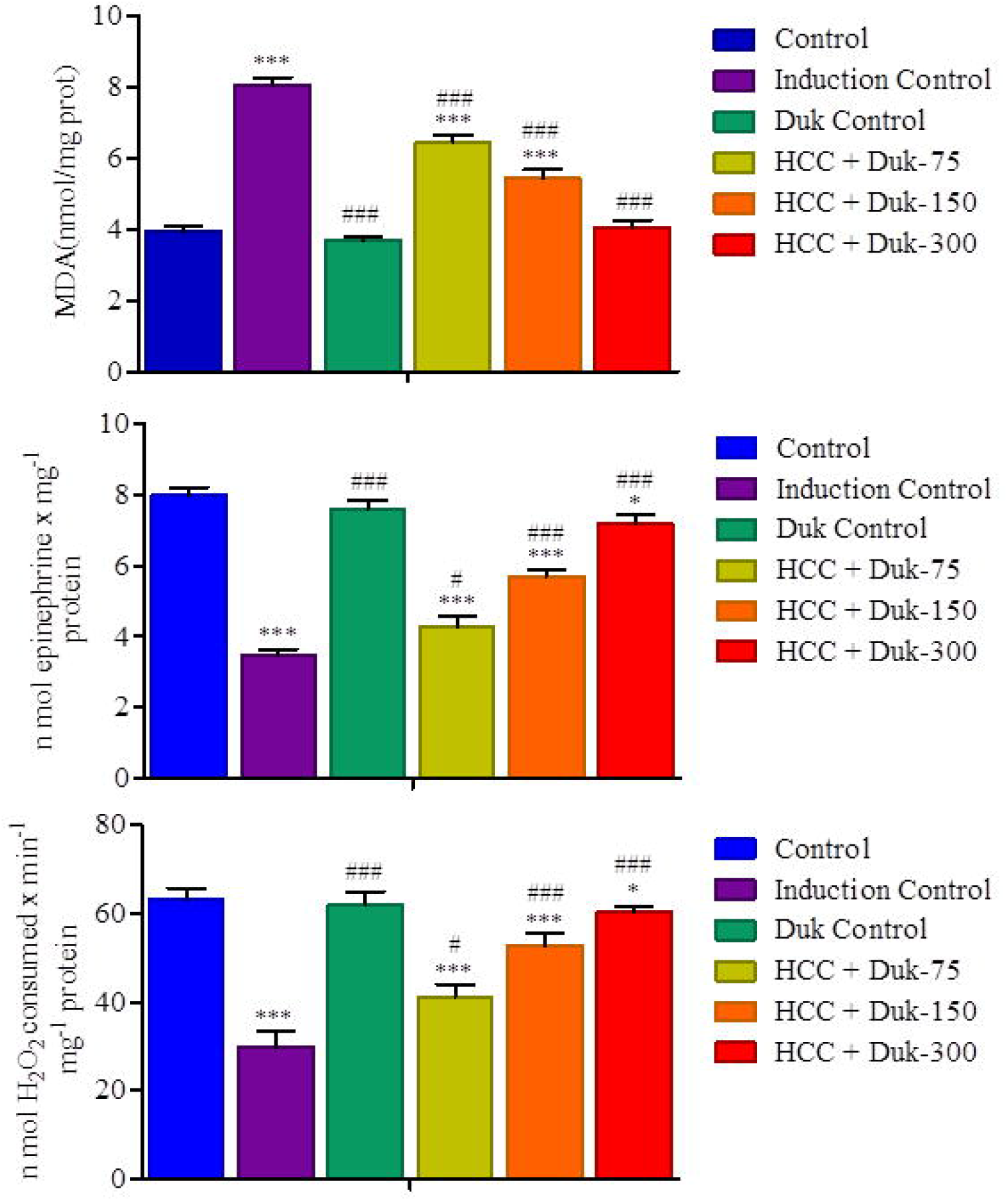
Effect of Duk on the hepatic levels of MDA, SOD and CAT in all experimental groups. Values are expressed as Mean ± SD (P<0.05 vs. control group*; P<0.05 vs. induction control group#). Abbreviations: HCC: Hepatocellular Carcinoma; Duk: Dawa-ulkurkum; HCC+Duk@75: Same dose of carcinogens as administered in induction control group + Duk at 75mg/kg dose; HCC+Duk@150: Same dose of carcinogens as ad-ministered in induction control group + Duk at 150mg/kg dose; HCC+Duk@300: Same dose of carcinogens as administered in induction control group + Duk at 300mg/kg dose; MDA: Malondialdehyde; SOD: Superoxide dismutase; CAT: Catalase

### 3.3. Reversal of liver architecture in HCC rats by Duk

The gross morphology of livers of the Duk only group was similar to the vehicle control group (Figure 4a and 4b). Histopathological examinations of the liver in the induction control group showed vacuolization of hepatocytes in the centrizonal area with the vari-tion of nuclear size. The cells in the nodule were large with clear cytoplasm and no si-nusoidal spaces. Clear nuclear atypia in the adenoma was seen (Figure 4c). However, these histopathological features reduced markedly in Duk-treated HCC rats in a dose-dependent manner (Figure 4d-4f). Our results validated that hepatotoxicity was accurately induced by DEN+ 2-AAF model. As a result of treatment with Duk, alterations in the liver microarchitecture disappeared in a concentration dependent manner. This ameliorative effect could be attributed to the antioxidant activity of Duk.

**Figure 4.**
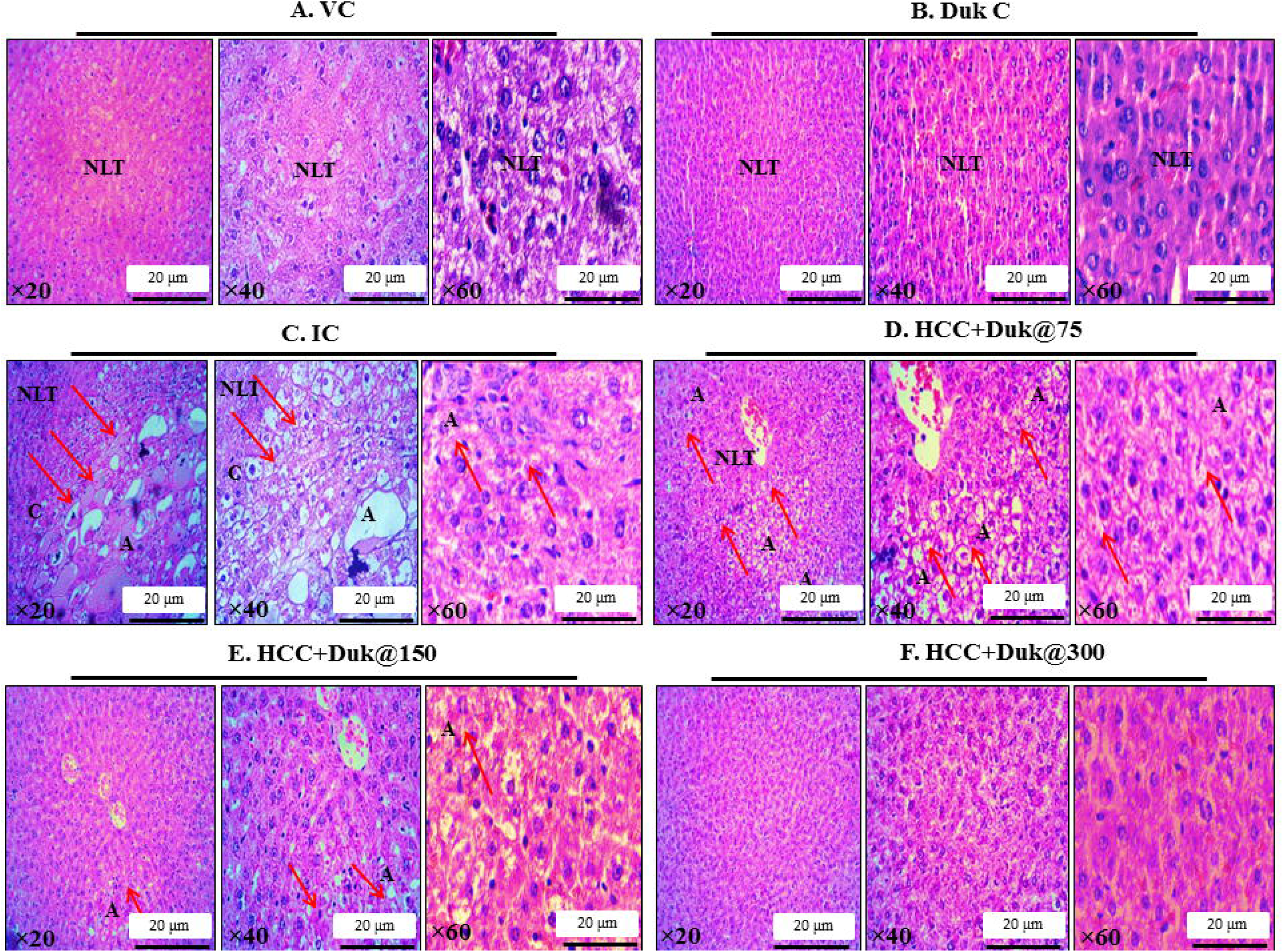
Photomicrographs showing histopathological changes in liver tissue: (a) Vehicle control group, showing normal liver microarchitecture (b) Duk control group, depicting well-organized liver cells into cell plates separated by sinusoids (c) Induction control group (DEN+2AAF; 16 weeks) presenting adenoma (Red arrows) in the parenchyma of liver section. Nuclear atypia and vacuolization in the cytoplasm can be seen (d) HCC+Duk@75 group, displaying multiple adenomas in the parenchyma of liver tissue. High-power photographs showing the clearing of cytoplasm and nuclear atypia (e) HCC+Duk@150 group, showing a reduced number of hepatic nodules and abnormal hepatocytes (f) HCC+Duk@300 group. Scale bar 20μm is given at all magnifications. Ab-breviations: VC: Vehicle Control; Duk C: Duk at 300mg/kg b.w. dose; IC: Induction Control; HCC: Hepatocellular Carcinoma, Duk: Dawa-ul-kurkum; HCC+Duk@75: Same dose of carcinogens as administered in IC group + Duk at 75mg/kg dose; HCC+Duk@150: Same dose of carcinogens as administered in IC group + Duk at 150mg/kg dose; HCC+Duk@300: Same dose of carcinogens as administered in IC group + Duk at 300mg/kg dose; NLT: Normal Liver Tissue, A: Adenoma, C: Capsule

To further confirm the activity of Duk against HCC, we immune-stained liver sections from the experimental animals for visualizing the presence of Bid, Caspase-9 and Cyclin D. The vehicle control and Duk only group showed very weak staining for Bid and Caspase-9 in a wide area. The staining intensity of Bid and Caspase-9 (Figure 5) in un-treated HCCs was slightly lower than that in the Duk-treated HCC groups. We observed deeper stain in sections of group 4 and 5 and lighter stain in sections of group 6. The de-crease in the expression levels of Bid and caspase-9 pro-apoptotic proteins in group 6 is possibly due to the protective effect of Duk at its higher dose against DEN + 2-AAF. In agreement, liver sections of rats in Group 6 (HCC + Duk @300mg/kg) showed a similar unaltered architecture of hepatocytes as found in vehicle control group. Further, the control and Duk-only groups also showed a weaker Cyclin D staining in a wide area than the induction control group, whereas HCC nodules showed a diffused and more intense staining pattern. The staining intensity of Cyclin D in untreated HCCs was slightly greater than that in the Duk-treated HCC groups and the effect was dose-dependent (Figure 5). We infer that Duk induces apoptosis by strong Bid and caspase-9 reaction in all nuclei of hepatocytes in the treatment group and protected cells from liver damage as shown by weaker reaction with Cyclin D.

**Figure 5.**
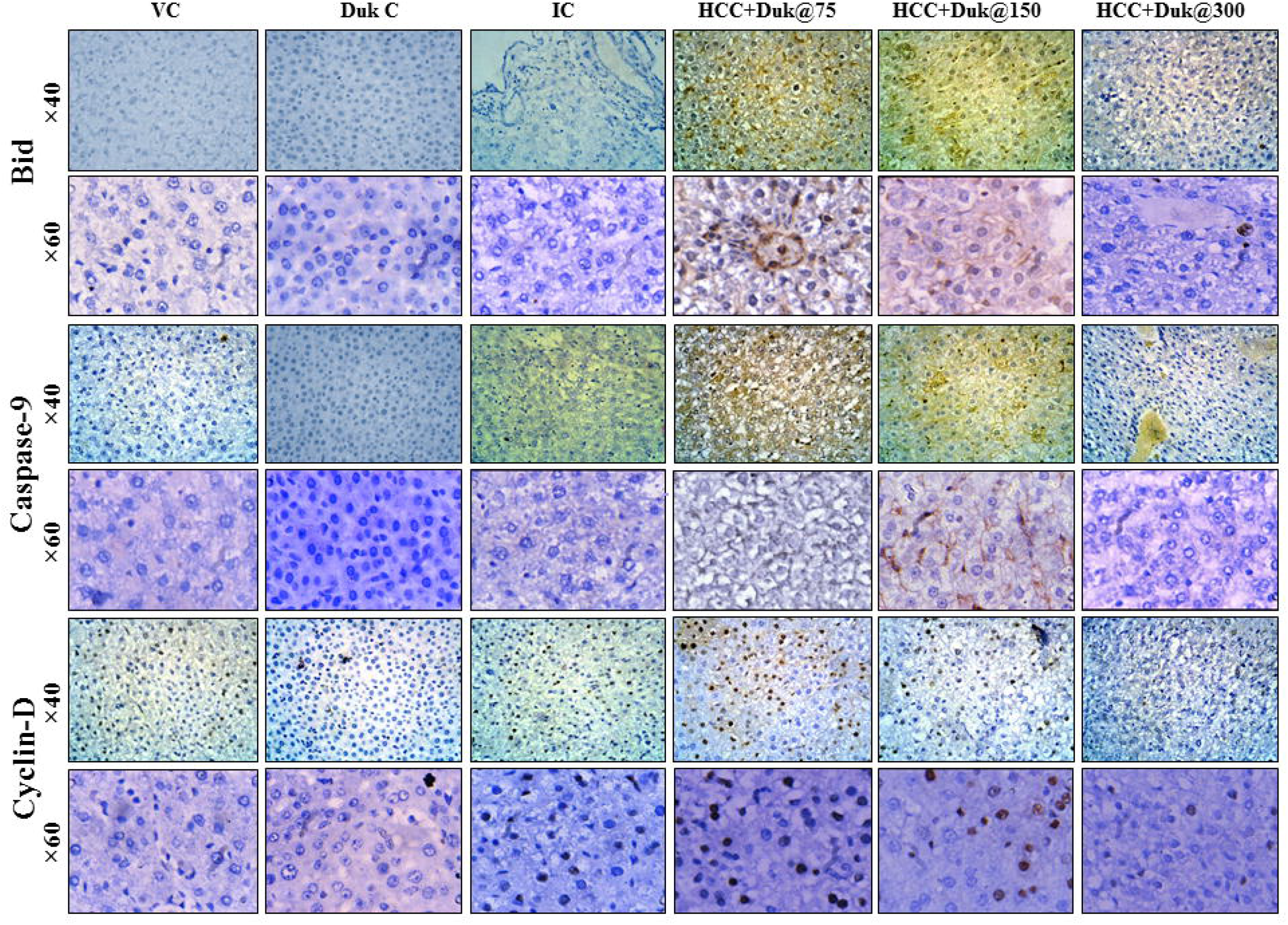
Expression of Bid, Caspase-9 and Cyclin D in rat liver sections as examined by immunohistochemistry. Abbreviations: VC: Vehicle Control; Duk C: Duk at 300mg/kg dose; IC: Induction Control; HCC: Hepatocellular Carcinoma, Duk: Dawa-ul-kurkum; HCC+Duk@75: Same dose of carcinogens as administered in IC group + Duk at 75mg/kg dose; HCC+Duk@150: Same dose of carcinogens as administered in IC group + Duk at 150mg/kg dose; HCC+Duk@300: Same dose of carcinogens as administered in IC group + Duk at 300mg/kg dose

## 4. Discussion

HCC is most commonly seen chronic liver disease caused by alcohol overuse, hepatitis B/C virus (HBV/HCV), or metabolic syndrome [22]. It is frequently multi-nodular when detected because of its synchronous progression. Natural preparations (as single herbs or polyherbal combinations) are gaining popularity in modern medicine due to their ease of use, low cost, safe usage, and various biological functions. The intricate interactions be-tween many phytoconstituents of polyherbal formulations aid in tumour cell targeting and provide a multi-dimensional approach to cancer management [23]. Duk is one such polyherbal Unani formulation prescribed for liver dysfunction, abdominal pain, anorexia and ascites. Previously, we have reported the organoleptic and phytochemical characteristics, antioxidant activity, and HPTLC fingerprint of Duk [10].

In the present study, a commonly used and widely accepted initiator of liver car-cinogenesis (DEN) was used. It gets activated by CYP450 system and leads to necrosis, hyperplasia and tumor. It is known to cause DNA mutation in normal liver cells and transform them into cancerous cells when challenged with another carcinogen (2-AAF) as a promoter. 2-Nitrosoflourine, a metabolite of 2-AAF is well known for disrupting the cell membrane, reacting with DNA and causing lipid peroxidation. Additionally, 2-AAF induces tumorigenesis by stimulating cell proliferation and inflammation [20].

Hepatocellular damage is associated with increased levels of serum hepatospecific enzymes (well established markers of liver function) [20]. In our study, administration of Duk has shown to significantly restore the levels of these serum markers. DEN+2-AAF have also been reported to generate lipid peroxidation products (MDA) and decrease the antioxidant enzymes [20]. It is linked with alterations in oxygen radical metabolism. We observed similar alterations in the induction control group and these effects were reversed by Duk in a dose dependent manner.

The biochemical findings were further supported by histopathological and immunohistochemical observations of the liver. The present study supported the fact that DEN+2-AAF initially induced damage around the central lobule, which further acted as precursor of cancer development. In the animals challenged with DEN, vacuolar cell starts appearing from the 5th week and gradually increase in number and size with time [24]. At the end of 16 weeks, we observed necrosis, adenomas formation and adenomatous nodules in the parenchyma cells of the livers of induction control group. These changes in the histopathological features of liver were attenuated after treatment with Duk, confirming the preventive effect of Duk.

Next, we selected three proteins for the immunohistochemistry: Bid, Caspase-9 and Cyclin D. Several studies reported that Cyclin D [25,26], Bid [27,28] and Caspase 9 [29,30] play important role in various stages of HCC. Cyclin D is well known for liver cell growth promotion and its overexpression is directly linked to HCC initiation. However, decreased expression of Bid and Caspase-9 is connected to the development and progression of HCC, showing their tumor suppressor role in HCC [25-30]. Our results gave a well-validated anticancer property for Duk, suggesting it could be complimentary a potent formulation against HCC and thereby necessitating further pre-clinical and clinical studies.

## 5. Conclusions

Although Duk is a well-established drug for various human ailments, our study reported for the first time its anti-cancer activity for liver cancer. However, more studies at the molecular and in vivo levels are necessary to understand the exact role and mechanism involved.

## Supporting information

Supplemental Figure 1. Seven herbal components of Duk

## Authors Contribution

Conceptualization & methodology: MG, HN, MS; Investigation: MG; Resources & Funding acquisition: MS; Writing—original draft preparation: MG, MS; Writing—review and editing: MG, MS; Supervision & project administration: MS. All authors agree to be accountable for all aspects of work ensuring integrity and accuracy. All authors have read and agreed to the published version of the manuscript.

## Institutional Review Board Statement

All the experimental procedures of this study were duly ratified by the Institutional Animal Care and Ethical Committee of Amity University, Noida, Uttar Pradesh, India (CPCSEA/IAEC/AIP/2018/05/06). The recommendations cited by the Committee for Control and Supervision of Experiments and Animals (CPCSEA) were used as guidelines for maintaining the animals during this period.

## Informed Consent Statement

Not applicable.

## Data Availability Statement

This manuscript contains most of the data. The corresponding author of this article, Maryam Sarwat, will provide further details upon reasonable request.

## Acknowledgments

None

