## Supplementary figures and images for "Pharmacological Studies on the Efficacy of Polyherbal Formulation Dawa ul kurkum in DEN-Induced 2-AAF-Promoted Hepatocellular Carcinoma in Male Wistar Rats"

### Supplemental Figure 1. Seven herbal components of Duk

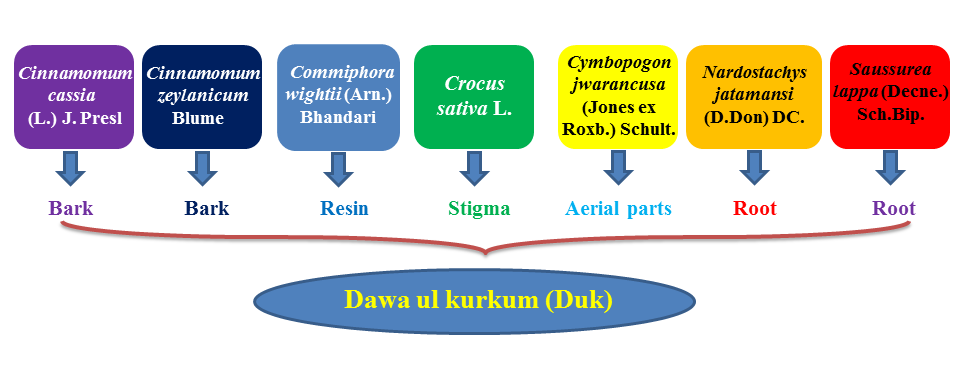
